# Continuous chlorophyll fluorescence measurements trace sudden cold spell effects on photosynthetic efficiency in a temperate mixed forest

**DOI:** 10.1101/2025.11.18.688898

**Authors:** Clara Stock, Stefanie Dumberger, Kathrin Kühnhammer, Markus Sulzer, Andreas Christen, Simon Haberstroh, Christiane Werner

## Abstract

Air temperature extremes and fluctuations are expected to become more frequent and have been shown to affect the physiological functioning of temperate forests across Europe. The exposure to sudden cold spells during summer combined with high light intensities can lead to photoinhibition of photosystem II (PSII) and thereby reduce photosynthetic efficiency. In this study, we aimed to analyse the dynamics of photoinhibition as well as related protection mechanisms of tall tree canopies during such cold spells. Therefore, we continuously assessed leaf level chlorophyll fluorescence (ChlF) and ecosystem carbon fluxes by eddy covariance in a temperate mixed forest in southern Germany during growing season 2024. While the deciduous, broadleaved *F. sylvatica* indicated chronic photoinhibition as a response to sudden cold spells, the evergreen, coniferous *P. menziesii* showed higher tolerance to low air temperatures. Both species increased thermal energy dissipation at PSII during cold spells indicating the activation of protective mechanisms. Likewise, both species exhibited mainly the sustained form of non-photochemical quenching (NPQ_s_) as a reaction to chilling temperatures and high light intensities which maintained elevated during the recovery phases. However, the dynamics of upregulation of photoprotection and recovery processes differed between the two tree species. Furthermore, the integration of ecosystem carbon exchange and continuous leaf level ChlF measurements gave valuable insights into the photosynthetic dynamics of the mixed forest canopy. This study emphasises the importance of the assessment of dynamic responses to future climate impacts on forest ecosystem on different scales.

## Introduction

The impact of extreme temperatures on temperate forest ecosystem functioning has lately been observed in several studies (Breshears et al. 2021; Marchin et al. 2022; Martini et al. 2022; Werner et al. 2025). Exposure to temperatures outside the species-specific optimum inhibits optimal photosynthetic functioning, especially when combined with high light intensities, even without the co-occurrence of edaphic drought conditions (Powles 1984; Huner et al. 1998; Öquist and Huner 2003). Therefore, chlorophyll fluorescence of photosystem II (PSII) is a sensitive tracer to monitor impacts on photosynthetic efficiency of high (Wittmann and Pfanz 2007; Haque et al. 2014; Tiwari et al. 2021; Húdoková et al. 2022; Martini et al. 2022) and low temperatures (Groom and Baker 1992; Solhaug and Haugen 1998; Oliveira and Peñuelas 2005; Wittmann and Pfanz 2007; Tyystjärvi 2008; Oivukkamäki et al. 2025) in combination with high light intensities. Yet, while many field studies are focusing on the effect of heatwaves on photosynthetic activity in forest ecosystems (Wohlfahrt et al. 2018; Martini et al. 2022; Gauthey et al. 2024), most of the studies investigating the effect of sudden cold temperatures on tree physiology have been limited either to seedlings in laboratory or greenhouse experiments (e.g. Oliveira and Peñuelas 2005; Robakowski 2005; Wittmann and Pfanz 2007), to late frost impacts (Ensminger et al. 2008; Linkosalo et al. 2014), or impacts of chilling air temperatures on evergreen species during winter (e.g. Martínez-Ferri et al. 2004; Porcar-Castell 2011; Porcar-Castell et al. 2012). However, although global air temperature is increasing (IPCC, 2023), both, cold and warm air temperature extremes, are likely to occur more frequently and with a higher intensity across Europe (Liu et al. 2025). Furthermore, extended growing seasons in Europe could cause departures during transitioning periods in spring and fall. Therefore, temperate forests in Central Europe will most likely not only face heat waves, but also chilling stress during the growing season, i.e. low (<15°C), but non-freezing temperatures (Hodgson et al. 1987; Theocharis et al. 2012). Rapid changes in air temperatures could have stronger impacts as there is no time for acclimation. However, how temperate forest species will respond physiologically to increased fluctuations and sudden changes of air temperature in the future and how it will affect photosynthetic functioning at the leaf and ecosystem scale is still uncertain. Continuous measurements of physiological parameters within forest canopies are therefore crucial for further research.

To assess a plants’ physiological state, chlorophyll fluorescence pulsed amplitude modulation (PAM) measurements have been used as a non-invasive and reliable technique for more than four decades (Schreiber et al. 1986; Krause and Weis 1991; Maxwell and Johnson 2000; Murchie and Lawson 2013). However, continuous, automated chlorophyll fluorescence (ChlF) measurements of dynamic responses of photosynthetic quantum use efficiency of PSII to sudden environmental changes, such as heat waves or cold spells, in tall forest canopies are still scars. Continuous *in situ* measurements could open new opportunities to capture the dynamics of photoinhibitory processes inside forest canopies, as well as energy partitioning at PSII and their impact on carbon uptake and therefore function as a sensitive indicator for the early onset of temperature-related changes in the photosystem (Porcar-Castell 2011; Magney et al. 2017; Meeker et al. 2021; Oivukkamäki et al. 2025). For instance, measurements of maximum quantum use efficiency (Fv/Fm) reflect the extreme sensitivity to cold temperatures of different tree species, such as silver birch (*Betula pendula*) (Oivukkamäki et al. 2025) or evergreen oak (*Quercus ilex*) (Oliveira and Peñuelas 2005).

Exposure to chilling temperatures induces a cascade of physiological responses, including the temperature-related decrease of the activity of enzymes, such as Rubisco, Rubisco-activase or ATP-ase, resulting in lower assimilation rates. Subsequently, light absorption on sunny days is excessive of what can be utilized for photochemistry and leads to overexcitation of PSII (Martínez-Ferri et al. 2004; Tyystjärvi 2008; Mathur et al. 2014; Verhoeven 2014). As a consequence, non-photochemical quenching (NPQ) is activated which leads to a change of the energy partitioning at PSII consisting in increasing thermal energy dissipation and decreasing effective quantum use efficiency (ΔF/Fm’) (Groom and Baker 1992; Long et al. 1994; Osmond and Grace 1995; Maxwell and Johnson 2000; Murchie and Ruban 2020). The reversible form of NPQ (NPQ_r_) protects PSII by enhancing the controlled loss of energy and recovers after dark adaptation (Ruban 2017). However, under the prolonged exposure to environmental stress, NPQ_r_ might not be able to fully protect PSII resulting in a diurnal decline of Fv/Fm (Ruban 2017). This dynamic photoinhibition is reversible and relaxes overnight (Werner et al. 2002). If the stress situation persists even longer, the sustained component of NPQ (NPQ_s_) gets activated (Demmig-Adams and Adams III 2006; Porcar-Castell 2011; Verhoeven 2014). NPQ_s_ is characterised by its overnight accumulation and is coupled to chronic photoinhibition, resulting in the sustained decline of predawn Fv/Fm values, when PSII is not able to fully recover overnight (Werner et al. 2002).

The susceptibility to photoinhibition and the capacity of photoprotection under high light intensities and chilling temperature, however, vary within different plant types. Evergreen tree species, for instance, generally show a higher capacity for non-photochemical quenching under high light intensity and low temperature and therefore more efficient protection mechanisms of PSII compared to deciduous tree species, which in the case of gymnosperms could partly be explained by additional electron sink pathways that help to prevent the over-reduction of the thylakoid membrane (Huner et al. 1998; Cavender-Bares et al. 2000; Ilík et al. 2017; Huang et al. 2021; Bag et al. 2023).

Moreover, combining continuous ChlF measurements in forest canopies with e.g. continuous gas exchange measurements can reveal deeper insights into the dynamics of photosynthesis (Magney et al. 2017; Meeker et al. 2021; Oivukkamäki et al. 2025; Sulzer et al. 2025). By combining these techniques, the ratio between electron transport rate (ETR) at PSII and leaf net carbon uptake (A_N_) can be calculated (ETR/A_N_). ETR/A_N_ serves as a tool to estimate a plant’s stress, but also supervise data quality of simultaneously assessed parameters (Perera-Castro and Flexas 2023). For instance, temperature-related stress can lead to an increase of ETR/A_N_, which indicates a higher use of electrons per fixed carbon and thus, the utilization of alternative electron sinks or pathways (Flexas et al. 1999; Dambrosio et al. 2006; Oivukkamäki et al. 2025).

In this study, we continuously assessed chlorophyll fluorescence at the ECOSENSE forest, a mixed forest canopy comprising mainly two temperate tree species, *Fagus sylvatica* (L.) and *Pseudotsuga menziesii* (MIRBEL) FRANCO, during the vegetation period of 2024 in the foothills of the Black Forest, Germany. Simultaneously, we recorded ecosystem carbon fluxes by Eddy covariance at a tall tower. Analogously to the parameter ETR/A_N_, we used the values of both techniques to calculate the parameter ETR/GPP to compare leaf level electron transport rate to whole ecosystem carbon uptake. Even though 2024 was the warmest year on record in Germany, there was no pronounced heat wave or drought period during the summer season, as it was a relatively wet year (DWD, yearly report 2024). However, there were several cold spells during late summer, both caused by low-pressure systems that directed cool air masses into Central Europe.

The overarching goal of this study was to investigate the dynamics of photoinhibition in a mature forest canopy based on high temporally resolved measurements of chlorophyll fluorescence and ecosystem carbon fluxes throughout the vegetation period 2024, with a particular focus on cold spells in late summer. We addressed three objectives: 1) understanding the underlying processes of photoinhibition and photoprotection in mature trees during late summer cold spells, 2) investigating differences between a broadleaved deciduous and an evergreen coniferous tree species and 3) analysing whether and how photoinhibition of PSII affects overall carbon fixation on an ecosystem scale and how continuous leaf level chlorophyll fluorescence measurements can enrich ecosystem carbon flux measurements to deepen our knowledge of photosynthetic dynamics in forest canopies during cold spells.

We hypothesize that the exposure to sudden cold spells at high light intensities late in the vegetation period decreases the photosynthetic efficiency and activity of the leaves by causing dynamic and chronic photoinhibition (*sensu* Werner et al. 2002). While we expect this effect to be stronger in the deciduous broadleaved species *Fagus sylvatica*, we assume higher tolerance and resilience of the evergreen conifer *Pseudotsuga menziesii*. We further postulate that the loss of photosynthetic efficiency during cold spells has a cascading effect on overall ecosystem carbon uptake.

## Material and Methods

### Study site and experimental design

The study was carried out in the ECOSENSE forest (Werner et al. 2024; Tesch et al. 2025) close to Ettenheim, Germany (48.2685 N, 7.8782 E), which is located at the foothills of the Black Forest. The forest is dominated by mature *F. sylvatica* and *P. menziesii* trees. On the field site, an eddy covariance tower (46 m) was installed in March 2024, allowing access to the sun crowns of both species (n = 3 per species) with platforms at ∼26 m height (Tesch et al. 2025). Studied *F. sylvatica* trees were on average 29.0 ± 1.6 m high with a mean diameter at breast height of 0.4 ± 0.1 m, studied *P. menziesii* individuals were on average 32.1 ± 3.8 m with a mean diameter at breast height of 0.4 ± 0.2 m. *F. sylvatica* and *P. menziesii* trees over the field site are between 55-110 and ∼45 years old, respectively. Mean annual air temperature and mean annual precipitation sum of the region measured by the closest official weather stations (station Lahr and station Ettenheim/Ettenheimmünster) from 1992 – 2020 were 11.0 °C and 911 mm, respectively (DWD, 2023). During the 2024 season, air temperature on the field site was measured every minute by a HygroVUE10 sensor (Campbell Scientific Inc., Logan, UT, USA) which was mounted at 27 m height on the tower scaffold in the crown space.

### Chlorophyll fluorescence

In total, three *F. sylvatica* and three *P. menziesii* trees, which were accessible from the tower platforms, were equipped with one MICRO-PAM measuring head each (Heinz Walz GmbH, Effeltrich, Germany) to measure active chlorophyll fluorescence continuously on plant leaves and needles, respectively. The measuring heads, controlled by a MONI-DA system (Heinz Walz GmbH, Effeltrich, Germany), recorded various chlorophyll fluorescence parameters (minimum and maximum fluorescence in the dark (F0, Fm) and in the light (Ft/Fm’), maximum (Fv/Fm) and effective quantum yield efficiency (ΔF/Fm’) of PSII and electron transport rate (ETR)), leaf-level photosynthetic photon flux density (PPFD) and leaf temperature at 15-minute intervals from 11^th^ of June to 15^th^ of November 2024. For the measurements, sun-exposed leaves were chosen. The cables of the measurement heads were attached to the branches with cable ties to distribute the weight of the device and prevent bending of the measured leaves. In case of *P. menziesii*, several needles were aligned next to each other and then fixed into the measuring head to ensure sufficient fluorescence signal. The measuring heads were kept on the same leaf (*F. sylvatica*) or branchlet (*P. menziesii*) as long as possible. However, especially during autumn, the measuring heads detached frequently due to heavy wind and precipitation events and in some cases the measured leaves or branches broke during storms. In these instances, the measuring head was re-installed as soon as possible to the same or a neighbouring leaf or branchlet with comparable sun exposure.

To initiate a chlorophyll fluorescence measurement, a saturation pulse was applied every 15 minutes using the internal clock modus of the MONI-DA. Measurement light was switched off between the measurements. The measurement light intensity was ∼1 µmol s^-1^ m^-2^ with a frequency of 15 Hz, while the saturation pulse had an intensity of ∼6,100 µmol s^-1^ m^-2^ and a width of 0.6 seconds. These settings were adjusted at the beginning of the measurement period by analysing the saturation pulse characteristics to ensure that saturation for both species in dark and light conditions was reached.

### Data processing

During the season, the internal clock of the MONI-DA system shifted slightly resulting in a time offset of about 4 min after three months. Timestamps were corrected during post-processing assuming the shift to be linear to be able to align the chlorophyll fluorescence measurements to other measured parameters such as air temperature and gross primary production (GPP).

Data were removed when measuring heads were detached abruptly from the branch or leaf, e.g., after heavy storms and when the fluorescence signal slowly decreased over time, indicating that the measuring head was slowly detaching from the leaf. Additionally, data was discarded when minimum fluorescence (F0 or Ft) was smaller than 50, as the signal to noise ratio gets too high, or when Fv/Fm or ΔF/Fm’ exceeded 0.84, which is considered unrealistically high (Björkman and Demmig 1987). Daily predawn values were calculated from the quality-controlled dataset by extracting the maximum Fv/Fm value during the night.

### Calculation of non-photochemical quenching

Total non-photochemical quenching (NPQ), sustained (NPQ_s_) and reversible non-photochemical quenching (NPQ_r_) were calculated using the following equations (1-3) according to Porcar-Castell (2011) and Zhang et al. (2025):

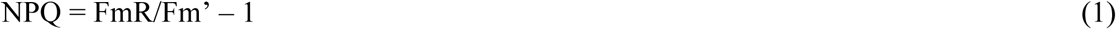

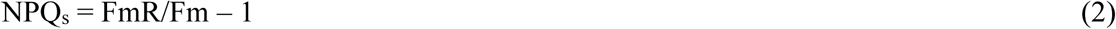

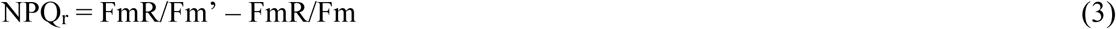

The required parameters for the equations were determined according to Porcar-Castell (2011): Maximum fluorescence (Fm) was assessed once per night, when Fv/Fm reached its maximum value. Fm’ refers to the maximum fluorescence from any given saturation pulse measurement. The reference value of maximum fluorescence (FmR) was determined during night, when Fv/Fm reached its maximum value and no downregulation was present. It has been proposed to determine one reference value (FmR) for a whole season, which is used to calculate all NPQ values (Porcar-Castell 2011). However, this requires a constant distance between the light guide and the measured area of the leaf (Zhang et al. 2025), which was not given in the present study for each tree throughout the entire season due to repeated detachment of the measuring heads during storms. Therefore, we identified periods per individual where a constant distance could be assured and extracted the FmR value from the maximum predawn Fv/Fm during the given period. This resulted in several FmR values per tree throughout the season which were then used to calculate non-photochemical quenching parameters for the respective periods. When NPQ values were slightly negative, probably due to movement of the measured branches, values were set to 0 as NPQ is considered a positive value.

### Gross primary productivity

The net CO_2_ exchange over the ECOSENSE forest was measured by eddy covariance (Aubinet et al. 2012) at 46 m above ground level, approximately 20 m above the forest canopy. For the eddy covariance measurements, we used a LI-7200RS enclosed CO_2_ and H_2_O infrared gas analyser (LI-COR Inc., Lincoln, NE, USA) and a CSAT3B three-dimensional sonic anemometer (Campbell Scientific Inc., Logan, UT, USA), both with a frequency of 20 Hz. To calculate the half-hourly fluxes from the 20 Hz measurements, EddyPro (version 7.0.9) was used (LI-COR Biosciences 2022). For further processing of the ecosystem fluxes REddyProc was applied (Wutzler et al. 2018), including u_*_-filtering (Papale et al. 2006), gap filling (Reichstein et al. 2005), and partitioning net ecosystem exchange (NEE) into gross primary production (GPP) and ecosystem respiration (R_eco_) using the daytime approach (Lasslop et al. 2010). To investigate from which tree species the measured net fluxes come from, flux footprint predictions were calculated for all available flux measurements of 2024, according to (Kljun et al. 2015) and superimposed with tree species and cover maps (details see (Moutahir et al. 2025).

### Definition of cold spells

Cold spells were defined as periods with diurnal and nocturnal maximum air temperature below 15°C and 12°C, respectively. Such conditions occurred during 12^th^ to 16^th^ of September (first cold spell) and during 02^nd^ to 05^th^ of October (second cold spell) and were caused by low-pressure systems that directed cool air masses into Central Europe. In between, there was one shorter cold spell (28^th^ of September – 30^th^ of September) but heavy wind and rain events led to detachment of all measuring heads and loss of data during this period. For the comparative analysis of the cold spells, data collected in preceding, intermediate and post cold spell periods, when air temperature was higher (diurnal >15°C, nocturnal >12°C), were also investigated (Table 1). The additional periods were selected to be as comparable as possible to the two cold periods in terms of the number of days and incident PPFD (Table 1).

**Table 1.**
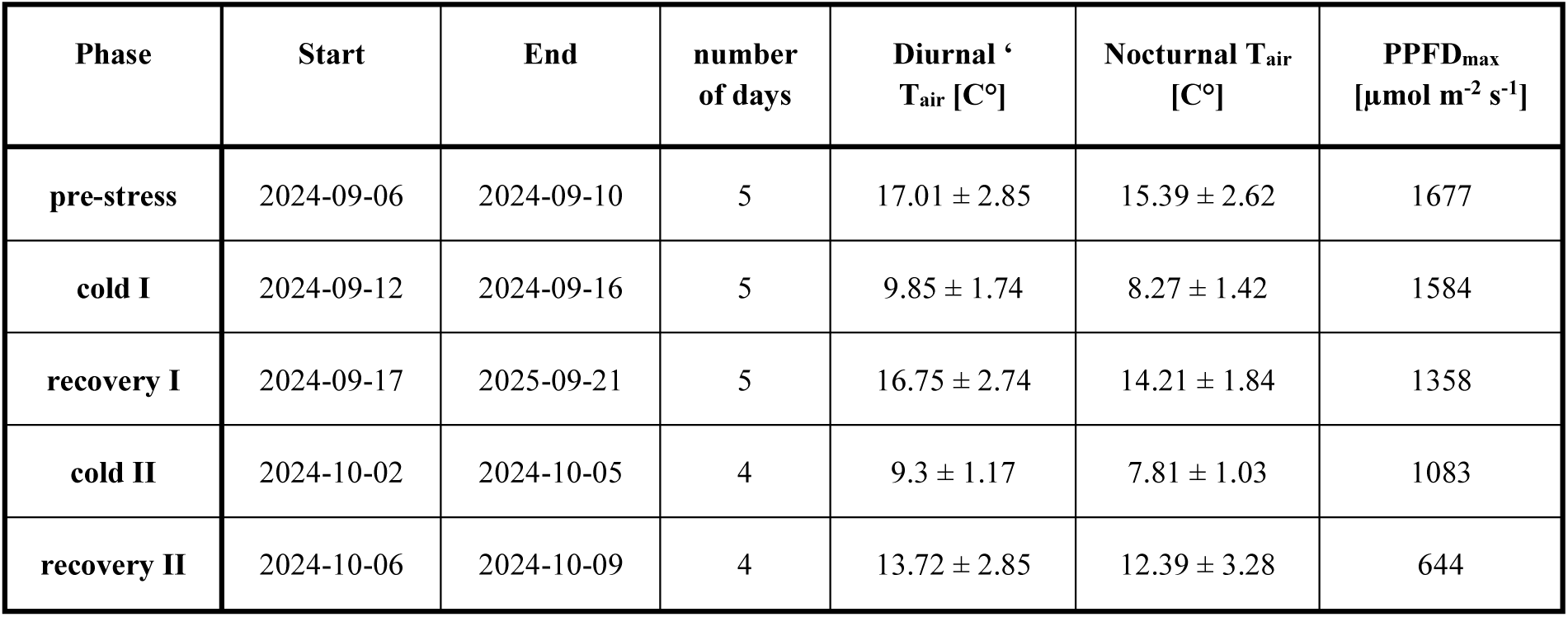
Characterization of the analysed time periods: start and end date, duration, mean diurnal and nocturnal air temperature in the canopy space (T_air_) with standard deviation and incident maximum photosynthetic photon flux density (PPFD_max_) acquired by the six MicroPAM measuring heads on the leaf level.

| Phase | Start | End | number of days | Diurnal $T_{air}$ [C°] | Nocturnal $T_{air}$ [C°] | $PPFD_{max}$ [$\mu mol\ m^{-2}\ s^{-1}$] |
| --- | --- | --- | --- | --- | --- | --- |
| pre-stress | 2024-09-06 | 2024-09-10 | 5 | $17.01 \pm 2.85$ | $15.39 \pm 2.62$ | 1677 |
| cold I | 2024-09-12 | 2024-09-16 | 5 | $9.85 \pm 1.74$ | $8.27 \pm 1.42$ | 1584 |
| recovery I | 2024-09-17 | 2025-09-21 | 5 | $16.75 \pm 2.74$ | $14.21 \pm 1.84$ | 1358 |
| cold II | 2024-10-02 | 2024-10-05 | 4 | $9.3 \pm 1.17$ | $7.81 \pm 1.03$ | 1083 |
| recovery II | 2024-10-06 | 2024-10-09 | 4 | $13.72 \pm 2.85$ | $12.39 \pm 3.28$ | 644 |

Due to wind and rain events measurement heads ripped off the leaf from time to time. Therefore, there were not always three replica per species throughout all analysed phases. However, in those cases the trends for the remaining two replicas were similar as evident from relatively low standard deviations.

### Statistical analysis

In order to evaluate species specific differences in daily predawn Fv/Fm values, a linear mixed effect model was calculated using the *lmer* function of the *lme4* package (Bates et al. 2015). Tree species and date were treated as fixed effects. Daily comparisons between species were calculated using the *emmeans* function of the *emmeans* package (Lenth et al. 2025).

To analyse air temperature and predawn Fv/Fm for *F. sylvatica* and *P. menziesii*, we first determined a linear mixed effects model per species using the function *lmer* from *lme4* package to verify whether there was a significant random effect caused by individual plants (Bates et al. 2015). Subsequently, we calculated a linear model using the *lm* function from base R. A breaking point was calculated using the function *segmented.lme* from the *segmented* package (Fasola et al. 2018). Finally, we calculated two simple linear models using the *lm* function, one before and one after the breaking point. For the calculation of the regression of *F. sylvatica*, values from 15^th^ of October onward were excluded, as the decrease of Fv/Fm during senescence in *F. sylvatica* might rather be related to structural changes in the leaf than to mere temperature effects.

To assure that model assumptions for linear and nonlinear mixed effect models were met, the normal distribution of residuals was visually validated. If model assumptions were violated, variables were transformed accordingly. Statistical significances were considered when p <0.05. All statistical analyses were performed using the open-source program R (version 4.5.1, R Core Team, 2025).

## Results

### Meteorological and carbon flux data

The year 2024 was relatively warm and wet, with a higher mean annual air temperature (12.4 °C) and total precipitation (1129 mm) compared to the long-term reference (1990 – 2020) (DWD). Daily mean air temperature from June to October 2024 fluctuated between 8 and 27°C (Fig. 1A). There were no pronounced heat waves, but distinct cold spells during late summer. During the measured period, incident radiation at leaf level reached maximum values of up to 2,007 µmol s^-1^ m^-2^ (Fig. 1B). Light intensities were relatively high with maximum values of 1,584 and 1,083 µmol s^-1^ m^-2^ during the first and the second cold spell, respectively (Fig. 1B, Table2).

**Figure 1:**
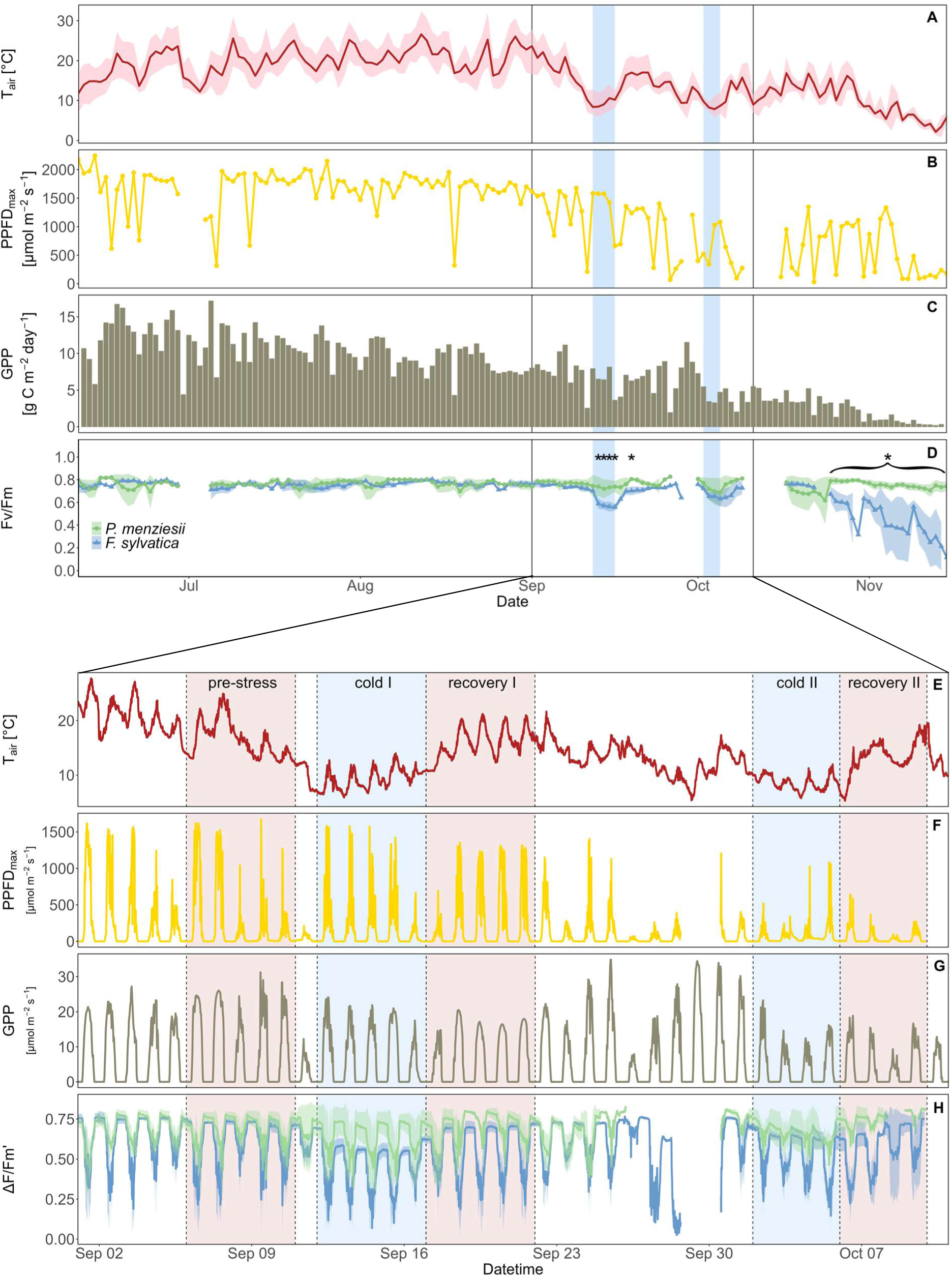
Daily average air temperature (T_air_) with daily maximum and daily minimum air temperature (A), leaf-level daily maximum photosynthetic active radiation (PPFD_max_) (B), daily sum of gross primary productivity (GPP) (C) and daily predawn Fv/Fm value per species (n=1-3) ± 1 SD (D). The lower part shows high resolution data from 1^st^ of September until 10^th^ of October: T_air_ recorded minutely (E), leaf-level PPFD_max_ value recorded every 15 minutes with the associated MicroPAM measurement (F), GPP recorded every 30 min (G) and mean effective quantum use efficiency (ΔF/Fm’) per species (n=1-3) ± 1 SD (H). Blue shaded areas (A-H) indicate the two cold spells (see Table 1), red shaded areas (E-H) the warmer phases (see Table 1), marked with vertical dashed lines. Asterisks indicate days with significant differences between the two species (p < 0.05).

Daily variability in GPP was mostly driven by incoming PPFD. There was no indication of stress periods in GPP over the vegetation period. At the end of the season when *F. sylvatica* started to senescence, GPP likewise dropped (Fig. 1C). We could not observe a temperature-related effect on GPP during the cold spells, as GPP did not show decreasing tendencies during the cold spells, but rather followed the incident light intensities, similarly to the warmer phases (Fig. 1C). Analysis of the flux footprint predictions showed that 65% of the available flux measurements in 2024 originated from areas covered by beech trees, 18% from areas dominated by Douglas fir and 17% from areas populated by other tree species.

### Temporal dynamics of chlorophyll fluorescence

Throughout the growing period, both species showed little variation and no significant differences in predawn Fv/Fm values, which varied between 0.7 and 0.82 (Fig. 1D). Average predawn Fv/Fm from 11^th^ of June to 11^th^ of September were 0.76 ± 0.02 and 0.77 ± 0.03 for *F. sylvatica* and *P. menziesii*, respectively. On the contrary, during two sudden cold spells in mid-September and beginning of October, predawn Fv/Fm values of *F. sylvatica* decreased abruptly to minimum values of 0.56 ± 0.03 and 0.63 ± 0.07 and remained low for four consecutive days during the first and the second cold spell, respectively (Fig. 1D, Table 2). After both cold spells, *F. sylvatica* predawn Fv/Fm values recovered immediately as soon as air temperatures started to rise again. During senescence, Fv/Fm values decreased constantly until the leaves were shed (Fig. 1D).

**Table 2.** Minimum and mean Fv/Fm, mean ΔF/Fm’, mean NPQ_s_, mean NPQ_r_ and mean ETR/GPP with standard deviation per species for all pre-defined phases (n=2-3). For the calculation of the means of ΔF/Fm’ and ETR/GPP only day values (PPFD > 0) were used. To determine the mean values per phase, we first calculated a mean (or minimum) value per individual per phase and subsequently the average per species and phase. Letters (a-c) indicate significant differences (p < 0.05) between the phases within one species and one parameter.

| Species | Phase | Fv/Fm<sub>min</sub> | Fv/Fm<sub>mean</sub> | $\Delta F/Fm'$<sub>mean</sub> | NPQ<sub>s</sub><sub>mean</sub> | NPQ<sub>r</sub><sub>mean</sub> | ETR/GPP<sub>mean</sub> |
| --- | --- | --- | --- | --- | --- | --- | --- |
| *F. sylvatica* | pre-stress | 0.72 ± 0.03<sup>a</sup> | 0.74 ± 0.02<sup>a</sup> | 0.52 ± 0.06<sup>a</sup> | 0.04 ± 0.02<sup>a</sup> | 0.51 ± 0.24<sup>a</sup> | 1.64 ± 0.90<sup>a</sup> |
|  | cold I | 0.55 ± 0.03<sup>b</sup> | 0.60 ± 0.03<sup>b</sup> | 0.36 ± 0.05<sup>a</sup> | 0.51 ± 0.19<sup>b</sup> | 0.94 ± 0.27<sup>a</sup> | 1.86 ± 0.92<sup>a</sup> |
|  | recovery I | 0.63 ± 0.02<sup>c</sup> | 0.69 ± 0.03<sup>a</sup> | 0.42 ± 0.06<sup>a</sup> | 0.28 ± 0.12<sup>ab</sup> | 0.78 ± 0.17<sup>a</sup> | 1.69 ± 0.96<sup>a</sup> |
|  | cold II | 0.63 ± 0.06<sup>bc</sup> | 0.67 ± 0.04<sup>ab</sup> | 0.47 ± 0.15<sup>a</sup> | 0.09 ± 0.08<sup>a</sup> | 0.51 ± 0.20<sup>a</sup> | 1.28 ± 0.51<sup>a</sup> |
|  | recovery II | 0.64 ± 0.09<sup>ac</sup> | 0.69 ± 0.07<sup>a</sup> | 0.54 ± 0.19<sup>a</sup> | 0.40 ± 0.02<sup>ab</sup> | 0.36 ± 0.13<sup>a</sup> | 1.08 ± 0.27<sup>a</sup> |
| *P. menziesii* | pre-stress | 0.75 ± 0.04<sup>a</sup> | 0.77 ± 0.03<sup>a</sup> | 0.64 ± 0.06<sup>a</sup> | 0.08 ± 0.03<sup>a</sup> | 0.42 ± 0.22<sup>a</sup> | 1.57 ± 1.11<sup>a</sup> |
|  | cold I | 0.72 ± 0.08<sup>a</sup> | 0.74 ± 0.08<sup>a</sup> | 0.58 ± 0.16<sup>a</sup> | 0.86 ± 0.89<sup>a</sup> | 0.46 ± 0.43<sup>a</sup> | 1.77 ± 1.82<sup>a</sup> |
|  | recovery I | 0.75 ± 0.06<sup>a</sup> | 0.78 ± 0.03<sup>a</sup> | 0.66 ± 0.06<sup>a</sup> | 0.77 ± 1.25<sup>a</sup> | 0.54 ± 0.54<sup>a</sup> | 1.43 ± 0.82<sup>a</sup> |
|  | cold II | 0.69 ± 0.12<sup>a</sup> | 0.72 ± 0.10<sup>a</sup> | 0.64 ± 0.07<sup>a</sup> | 0.58 ± 0.53<sup>a</sup> | 0.48 ± 0.14<sup>a</sup> | 1.54 ± 0.40<sup>a</sup> |
|  | recovery II | 0.76 ± 0.08<sup>a</sup> | 0.79 ± 0.04<sup>a</sup> | 0.71 ± 0.01<sup>a</sup> | 0.32 ± 0.45<sup>a</sup> | 0.31 ± 0.11<sup>a</sup> | 1.16 ± 0.14<sup>a</sup> |

In contrast to *F. sylvatica*, mean predawn Fv/Fm values of *P. menziesii* were not strongly affected by the first cold spell and remained above 0.7 (Fig 1D, Table 2), resulting in significant differences between both species (p < 0.05) (Fig. 1D). Throughout the second cold spell, *P. menziesii* mean predawn Fv/Fm also decreased slightly to 0.689 on 5^th^ of October. Hereafter, simultaneous to *F. sylvatica*, the values also recovered quickly as soon as air temperature increased. Significant differences between both species emerged during senescence from 25^th^ of October onwards as Fv/Fm values of *F. sylvatica* decreased continuously (p < 0.05) (Fig. 1D). Despite being an evergreen species, also *P. menziesii* predawn Fv/Fm values concurrently decreased slightly during this phase (Fig 1D).

High time-resolution data (Fig. 1E-H) indicated that only the combination of cold air temperature and high light exposure induced the reduction of quantum use efficiency: The sudden air temperature decline from minimum nocturnal values of 11.6 °C (10^th^/11^th^ of September) to minimum nocturnal values of 6.5 °C(11^th^/12^th^ of September) at the beginning of the first cold phase did not have an immediate effect on the nocturnal Fv/Fm values of both species, which kept relatively stable during the night of 11^th^/12^th^ of September (Fig. 1H). Only in combination with the high light intensities of the following day (12^th^ of September), ΔF/Fm’ decreased significantly in *F. sylvatica* during the day in comparison to the previous period with comparable PPFD values (Fig. 1F) and did thereafter not fully recover during the following night (Fig. 1H). The last day during the first cold spell and the following day showed lower incident PPFD values, which immediately resulted in higher ΔF/Fm’ during the day and higher Fv/Fm values during the subsequent nights (Fig. 1F and H).

### Dependency of predawn Fv/Fm on air temperature

The calculated breaking point of the relation between predawn Fv/Fm values of *F. sylvatica* and air temperatures during the whole growing season was at 16.5°C. While predawn Fv/Fm reached maximum values and remained stable above this breaking point, they started to decline with decreasing air temperatures below the breaking point (Fig. 2A). No heat-induced decline in Fv/Fm could be observed over the summer months, however, highest air temperatures in 2024 were moderate with a max of 32.5 °C. For *P. menziesii*, there was no significant effect of air temperature on predawn Fv/Fm. Nonetheless, we observed a weak tendency of predawn Fv/Fm values to decline below 10°C in *P. menziesii* (Figure 2B), but values were generally less affected by low air temperature than in *F. sylvatica*. The low temperature effect on Fv/Fm was not solely a seasonal effect, as full recovery was observed later in the season (Fig. 2).

**Figure 2:**
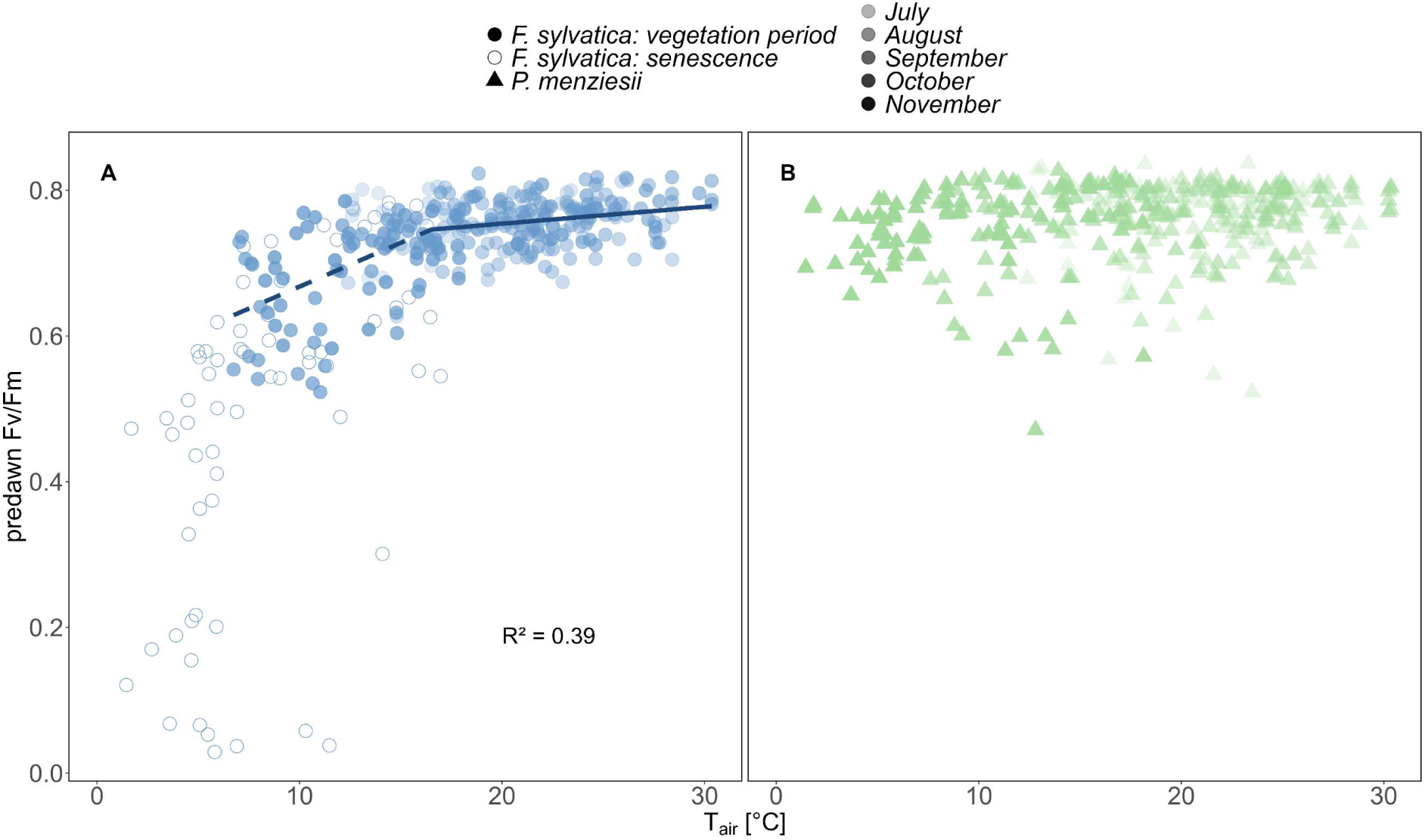
Relationship between air temperature and daily predawn Fv/Fm value for F. sylvatica (A) and P. menziesii (B) with regression lines (mixed effect models) for F. sylvatica values before (dashed; y = 0.55 + 0.012x) and after (solid; y = 0.71 + 0.002x) the calculated breaking point (16.49°C). The colour code indicates the time during the season. Dots (F. sylvatica) and triangles (P. menziesii) represent one predawn value per individuum and day. Open symbols of F. sylvatica visualize measurements during senescence (after 15^th^ of October) and were not considered in the mixed-effect model.

### Comparison of cold and warm periods

To enable a deeper analysis of the diurnal dynamics of the different ChlF parameters during the defined cold and recovery phases (Table 1), we calculated mean diurnal courses for PPFD, ΔF/Fm’, NPQ and ETR/GPP for each of the phases (Fig. 3). Both species were measured at the same canopy height (26m), however the measured leaves of *F. sylvatica* were exposed to higher mean incident light intensities than the measured needles of *P. menziesii* throughout the day, due to differences in canopy structure and leaf/needle orientation (Fig. 3A). The differences in incident light were more pronounced during the sunnier phases (pre-stress, cold I and recovery I) and decreased during the cloudier phases (cold II and recovery II), where the incident PPFD of the two species only differed slightly (Fig. 3A).

**Figure 3:**
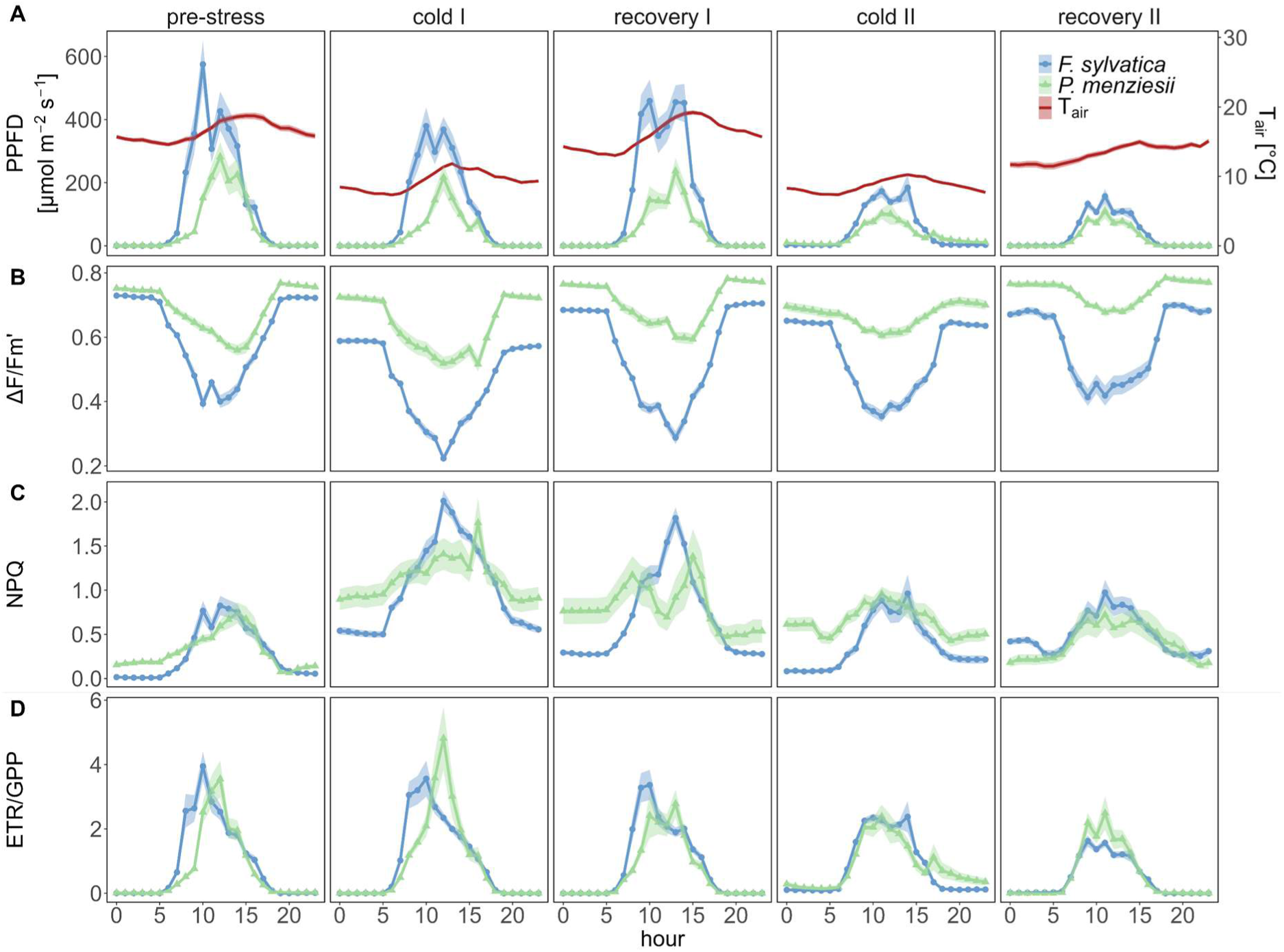
Mean diurnal courses during the five defined phases (Table 1): hourly mean values of PPFD and air temperature (A), ΔF/Fm’ (B), total regulated thermal energy dissipation (NPQ) (C) and ETR/GPP (D) per species (n=1-3). Shaded areas (A-D) indicate standard error. Note that for the calculation of ETR/GPP the same GPP values were used for each species.

Diurnal dynamics of ΔF/Fm’, NPQ and ETR/GPP were mostly driven by light dynamics following the diurnal courses of PPFD tightly (Fig. 3). Both species showed a decline of ΔF/Fm’ values during the cold spells, which was more pronounced in *F. sylvatica*, but still visible in *P. menziesii* (Fig. 3B). This decrease of quantum use efficiency during the cold spells was accompanied by a simultaneous increase of NPQ in both species (Fig. 3C). Interestingly, this temperature-induced increase of NPQ was of comparable magnitude in both species. Given that *P. menziesii* exhibited a weaker reaction in ΔF/Fm’ compared to *F. sylvatica*, the comparable increases in NPQ result in species-specific different ratios between ΔF/Fm’ decrease and NPQ increase during the cold spells (Fig. 3B & C). For the comparison of leaf level electron transport rate to whole ecosystem carbon uptake we calculated the parameter ETR/GPP. The response of ETR/GPP values to the cold spells was weak, but we could still observe a slight increasing tendency during the cold spells, taking into account total PPFD values being slightly lower than in the respective pre-stress phases (Fig. 3A & D, Table 2). In general, all parameters reacted stronger during the first and less pronounced during the second cold spell, which was characterized by lower incident light levels (Fig. 3).

Dynamics of ΔF/Fm’ during recovery differed between the two species. *F. sylvatica* showed increased ΔF/Fm’ during the recovery phases compared to the previous cold spells, but still decreased levels compared to the pre-stress phase (Fig. 3B). *P. menziesii*, on the other hand, recovered quickly reaching similar ΔF/Fm’ values in relation to PPFD as during pre-stress. NPQ values kept remarkedly high during the first recovery phase in both species (Fig. 3C).

### Sustained (NPQ_s_) and reversible (NPQ_r_) thermal energy dissipation

In order to gain a more mechanistic insight into the species-specific photoprotective dynamics, we separated the protective responses into reversible (NPQ_r_) and sustained non-photochemical quenching (NPQ_s_) (Fig. 4). While both species increased NPQ_s_ as a reaction to the cold spells, the proportion of the two forms of NPQ varied between the species. *P. menziesii* showed a higher portion of sustained NPQ during the first cold spell, while *F. sylvatica* increased both – sustained and reversible NPQ. Total NPQ_s_ values were higher in *P. menziesii* during the first cold spell, but recovered quicker than in *F. sylvatica*, reaching full recovery after five days. *F. sylvatica,* contrastingly, maintained elevated NPQ_s_ values over nine days after the first cold spell and no full recovery could be observed (Fig. 4). During the second cold spell, both species increased NPQ_s_ levels again. However, this time *F. sylvatica* showed a delayed increase, but enhanced values during the beginning of the second recovery phase (Fig. 4).

**Figure 4:**
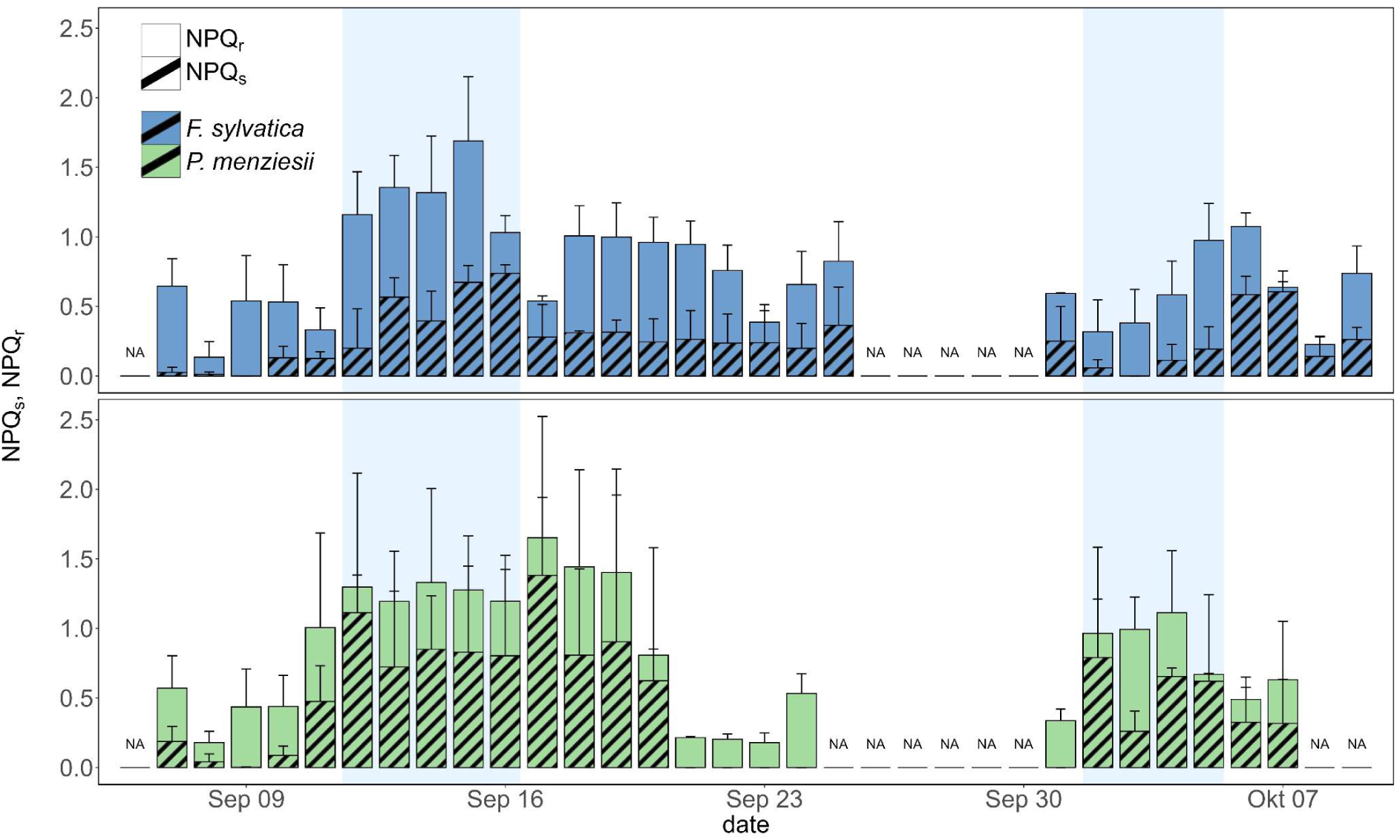
Daily mean sustained and reversible thermal energy dissipation: NPQ_s_ + 1 SD (hatched part of the bars) and NPQ_r_ + 1 SD (solid part of the bars) of F. sylvatica (upper panel) and P. menziesii (lower panel) from 06^th^ of September to 9^th^ of October. Blue shaded areas indicate analysed cold spells.

## Discussion

The cold spells during late summer 2024 caused a substantial species-specific decline in effective light use efficiency and clear photoinhibitory effects, particularly in *F. sylvatica*. Moreover, they induced a significant increase in sustained and reversible non-photochemical quenching (NPQ_s_). Nevertheless, ecosystem carbon fluxes were not affected indicating that ecosystem-scale carbon fixation rates remained stable during the cold spells despite the decrease in leaf-scale photosynthetic efficiency.

### Significant effect of chilling temperatures and high light intensities on photosynthetic efficiency

During the day, in light adapted plants the simultaneous exposure to low air temperature and high light intensities leads to an increase in thermal energy dissipation (NPQ) which is coupled to a decline in ΔF/Fm’ (Dambrosio et al. 2006; Busch et al. 2009; Porcar-Castell 2011). This change of energy partitioning at PSII serves as a protection mechanism that aims to prevent the over-excitation of PSII and could be shown in the present study during two summer cold spells in the canopy of mature *F. sylvatica* and *P. menziesii* trees (Fig. 1H, Fig. 3, Fig. 4). Both species responded with stronger decreases of ΔF/Fm’ and stronger increases of NPQ during the first than during the second cold spell (Fig. 3B, C), despite air temperature being even lower during the second cold spell (Table 1). These different dynamics were caused by generally higher light intensities during the first cold spell, which is consistent with the results of other studies and emphasises that only the co-occurrence of chilling temperatures with high light-intensities has a significant negative effect on the efficiency of PSII (Hodgson et al. 1987; Tyystjärvi 2008). The continuous assessment of ChlF parameters in this study enabled not only the analysis of general NPQ dynamics during summer cold spells, but also the separation into the reversible and sustained forms of NPQ. In our study, the prolonged exposure to chilling temperatures and high light intensities induced chronic photoinhibition (Werner et al. 2002) as indicated by the sustained down-regulation of predawn Fv/Fm values (Fig. 1D, H, Table 2). Simultaneously, the sustained form of NPQ accumulated over night in both species and even continued elevated as predawn Fv/Fm already recovered (Fig. 4). Previous studies have shown that the reversible and the sustained form of NPQ act in an additive way, when NPQ_r_ alone is insufficient to cope with the ongoing stress (Porcar-Castell 2011; Verhoeven 2014). This could be confirmed in the present study, as NPQ_r_ did not decrease during the increase of NPQ_s_ due to the cold stress and even increased slightly in *F. sylvatica*. This indicates that additionally to the dynamic, diurnal upregulation of reversible NPQ during the day, plants increase sustained NPQ when dynamic protection becomes insufficient.

### Stress reactions and recovery dynamics differ between broadleaved deciduous and evergreen coniferous species

We observed a higher susceptibility of Fv/Fm to chilling temperatures in *F. sylvatica* compared to *P. menziesii* (Fig. 2)*. F. sylvatica* showed a relatively strong decline of predawn Fv/Fm in response to chilling air temperatures and high light intensities during the preceding day. This is in line with a meta study by Neri et al. (2024) who analysed the influence of air temperature on maximum quantum use efficiency in different plant functional types (PFT) showing that, broadleaf deciduous trees were classified as trees with strong cold and hot resilience but a rather small range of tolerance, while coniferous evergreen temperate trees, were defined as plants with a wider temperature tolerance range (Neri et al. 2024). The sensitive response of Fv/Fm to chilling temperatures of broadleaved deciduous tree species has also been observed in other studies, comprising potted, as well as mature trees (Cavender-Bares et al. 2000; Oivukkamäki et al. 2025). Interestingly, *P. menziesii* – despite being only slightly affected in ΔF/Fm’ during the first cold spell compared to *F. sylvatica* and despite maintaining the capacity to fully recover predawn Fv/Fm values overnight – still increased NPQ levels during the first cold spell to a comparable level as those of *F. sylvatica* (Fig. 3C). Furthermore, *P. menziesii* showed a higher proportion of sustained NPQ, while *F. sylvativa* increased both, sustained and reversible NPQ, during the cold spells. This indicates that while the upregulation of NPQ in *F. sylvatica* could not fully prevent the temporal reduction in quantum use efficiency of PSII, *P. menziesii* was able to protect and maintain photosynthetic efficiency by upregulating thermal energy dissipation. We acknowledge the slightly different light levels the two species were exposed to due to their canopy structure and positioning of branches, which led to different incident PPFD intensities between the species (Fig. 3A). However, the pronounced increase of NPQ in *P. menziesii* during the first cold spell demonstrates that even the lower light levels in *P. menziesii* were sufficient to induce a temperature- and light-dependent reaction of the photosynthetic apparatus. Further species-specific differences in NPQ dynamics emerged during the recovery phase after the first cold spell (Fig. 4). While NPQ_s_ took four days in *P. menziesii* until total recovery, we could not observe a recovery even after nine days in *F. sylvatica*. The relaxation of sustained NPQ happened relatively quick in *P. menziesii*, but took longer in *F. sylvatica*, indicating a more flexible protection in our coniferous species vs. a sustained down-regulation in our broadleaved species. In general, evergreen species are known to have a higher capacity of upregulating thermal energy dissipation under the exposure to light and cold temperatures compared to deciduous species, which is considered an adaptation to a greater need for photoprotection during winter season and might be related to a larger violaxanthin pool in the dark in evergreen species compared to broadleaved deciduous species (Ebbert et al. 2005; Demmig-Adams and Adams III 2006; Huang et al. 2021). Additionally, gymnosperms, have been shown to own an additional alternative electron pathway, called the Flavodiiron-dependent pathway, that enables the quick removal of electrons from photosystem I (PSI) in a Mehler-like reaction and therefore, ensures a rapid oxidation of PSI (Ilík et al. 2017; Bag et al. 2023). This does not only help to protect PSI in the case of an over-excitation, but contributes in maintaining the electron transport form PSII to PSI (Ilík et al. 2017; Bag et al. 2023). It is assumed that angiosperms lack this mechanism, which could be an additional explanation for the different reactions of *F. sylvatica* and *P. menziesii* to the cold spells, where *P. menziesii* could maintain the electron transport more efficiently than *F. sylvatica*.

### Short term cold spells do not affect ecosystem productivity in late summer

ETR/GPP values in our study mainly followed the diurnal dynamics of PPFD (Fig. 3 A & D) which was consistent with the diurnal patterns of ETR/A_N_ shown by Oivukkamäki et al. (2025). We observed a weak tendency of higher ETR/GPP values during the cold spells compared to the other phases (Fig. 3A & D, Table 2). This indicates a higher usage of electrons per fixated carbon and could be explained by the activation of alternative electron sinks or pathways as a reaction to chilling stress (Perera-Castro and Flexas 2023). However, at ecosystem level, against our expectations, there was no measurable impact of the cold spells on overall GPP, although the quantum use efficiency on the leaf level declined, particularly in *F. sylvatica* (Fig. 1). This indicates that the impaired photosynthetic efficiency observed on the leaf level was not (yet) strong enough to impact total ecosystem fluxes. This could partly be due to the contribution of the shade crown. In our study, we only assessed ChlF parameters in sun exposed leaves. However, shade leaves have been shown to contribute significantly to overall GPP (He et al. 2018; Chen et al. 2020) and might therefore have a buffering effect during photoinhibition that affects sun leaves stronger than shade leaves.

Previous research has shown that chlorophyll fluorescence parameters are sensitive indicators of the early onset of photoinhibition – often preceding detectable changes in photosynthetic performance – as demonstrated in *F. sylvatica* and *Fraxinus excelsior* seedlings (e.g. Einhorn et al. 2004). Our findings extend this understanding to mature trees, showing that fluorescence-derived parameters may serve as early stress indicators and as a valuable addition to carbon flux measurements. Moreover, additional parameters, such continuous leaf level gas exchange or remotely sensed passive chlorophyll fluorescence, may aid to integrate responses across scales. This highlights the value of combining scale-integrated approaches of continuously assessed parameters in tree canopies to detect early stress responses (Werner et al. 2021; 2024).

## Conclusion

In conclusion, we found a significant impact of sudden cold spells on the photosynthetic efficiency at the end of the summer, when radiation levels were still high, particularly in a broadleaved but to a lower extend also in a coniferous species. Both species responded to the stress exposure with an increase in photoprotective mechanisms, in form of sustained NPQ. *P. menziesii* showed a higher portion of sustained NPQ, but also recovered quicker than *F. sylvatica* when temperatures increased again. While ecosystem carbon uptake was not (yet) affected by the cold spells, we clearly demonstrate that continuous chlorophyll fluorescence measurements can be used as a valuable and immediate parameter for early stress detection in mature forest trees.

## Acknowledgements

We acknowledge funding from the DFG in the project SFB1537 (ECOSENSE, project ID: 459819582). We gratefully acknowledge permission from the city of Ettenheim to set-up our field site in their city forest. Furthermore, we want to thank Delon Wagner and Josef Strack for their technical support on the field site.

## Data Availability

The data that support the findings of this study are available from the corresponding author upon reasonable request.

